# The influence of virtual visual stimulus amplitude to induce standing postural responses

**DOI:** 10.64898/2026.06.30.735222

**Authors:** Baptiste Toussaint--Malard, Frédéric R. Danion, Charlotte Le Mouel, Arnaud Decatoire, Pierre Laguillaumie, Maxime Billot, Romain Tisserand

## Abstract

Upright postural control during movement relies on multisensory integration. Yet, the frequency-specific contribution of vision remains poorly characterized in virtual reality (VR). This study investigated how multi-sine visual stimulation amplitude delivered in VR influences standing postural responses. Fifteen healthy adults stood on a force plate wearing a VR headset. Visuo-postural coupling was assessed through coherence and gain analyses between a multi-sine signal (10 sinusoids, 0.12 to 1 Hz) oscillating a virtual environment in one of four amplitudes (0.5, 1, 2, 4 degrees peak-to-peak) and the anteroposterior whole-body angle. All amplitudes elicited measurable postural responses. Increasing amplitude significantly increased postural oscillation and tended to increase coherence, while gain significantly decreased. These results are consistent with a nonlinear control system. The 2 degrees amplitude elicited the largest gain with significant coherence across all stimulated frequencies, suggesting it is suitable for studying visual contributions to postural control during movement execution.

## 1. Introduction

Upright postural control relies on the integration of visual, proprioceptive, and vestibular inputs. Vision plays a key role in movement planning, yet its contribution to whole-body movement execution remains unclear. Visual stimulation (VS) from a large oscillating screen can elicit postural responses in standing humans (Peterka, 2002), with similar results observed using virtual reality (VR) headsets (Assländer et al., 2023). Postural responses are characterized by coherence (linear input-output relationship across frequencies) and gain (output amplitude relative to input), both peaking near 0.5 Hz and progressively decreasing at higher frequencies. Gain additionally exhibits an amplitude-dependent profile, diminishing as stimulus amplitude increases. Previous studies have relied on sinusoidal or pseudo-random ternary sequences (PRTS) stimulations. Yet, single sinusoids induce habituation over time (Ketterer et al., 2022), while PRTS provide broadband excitation that limits the isolation of frequency-specific responses (Peterka, 2002). Consequently, the investigation of frequency-specific visual contributions to postural control in VR remains limited. Stochastic stimulations, used for vestibular inputs, enable continuous spectral sampling without habituation (Dakin et al., 2007). Inspired by this approach, an unpublished preliminary study from our group showed that multi-sine VS in VR elicited postural responses without habituation, but did not identify the impact of stimulus amplitude. The present study aimed to fill this gap, hypothesizing that large amplitude maximizes coherence and that lower amplitude maximizes gain.

## 2. Methods

### 2.1 Participants

Fifteen healthy adults (4 women, 26.7 ± 8.6 years old; 177.1 ± 10.9 cm; 73.8 ± 13 kg) participated. The recruitment and testing procedures complied with the Declaration of Helsinki and were approved by the University Ethics Committee.

### 2.2 Experimental design

Participants stood upright on a forceplate (40×60 cm) with feet hip-width apart and arms alongside the body for all 90s trials. They first completed two trials without VR (eyes open, EO, and eyes closed, EC), followed by one trial wearing a VR headset (HTC VIVE Focus Vision, 120 Hz) viewing a static virtual environment (“static” condition). Subsequently, the same environment oscillated back and forth around the mediolateral ankle axis following a multi-sine signal (sum of 10 sinusoids: 0.12, 0.2, 0.29, 0.37, 0.44, 0.5, 0.59, 0.68, 0.8, and 1 Hz) with fixed amplitudes and phases (“dynamic” conditions). Four peak-to-peak amplitudes were tested (0.5°, 1°, 2°, 4°), each repeated three times, yielding 12 randomly presented dynamic trials.

### 2.3 Data Analysis

The center of pressure trajectory, extracted from forceplate data (2000 Hz), was used to compute the anteroposterior (AP) body angle via an inverted pendulum model (Winter, 1998). The VS–postural response relationship was assessed using FFT, coherence, and gain analyses, with the EO condition power spectrum subtracted from other conditions to isolate stimulus-induced responses. For each condition, 65.56 seconds per trial were extracted (2^17^samples), concatenated first within participants and then across participants into a single pooled array enabling coherence computation at an acceptable significance level (Dakin et al., 2007). FFT, coherence, and gain values were extracted at stimulation frequencies, averaged across frequencies (Table 1), and compared using a paired sign test.

**Table 1.** Mean FFT (10-3), coherence and gain between VS and AP body angle averaged across VS frequencies in each condition. Values are reported as mean ± SD across stimulated frequencies. Cond indicates Conditions. Credit: Baptiste Toussaint--Malard

| Cond | FFT ($10^{-3}$) | Coh | Gain ($^{\circ}/^{\circ}$) |
| --- | --- | --- | --- |
| EO | $0 \pm 0^*$ | / | / |
| EC | $5 \pm 4^*$ | / | / |
| Static | $1 \pm 5^*$ | / | / |
| 0.5° | $8 \pm 5^*$ | $0.15 \pm 0.11$ | $1.25 \pm 0.20^{\beta}$ |
| 1° | $10 \pm 3^*$ | $0.17 \pm 0.11$ | $0.71 \pm 0.07^{\alpha}$ |
| 2° | $15 \pm 4^*$ | $0.19 \pm 0.10$ | $0.39 \pm 0.04^{\mu}$ |
| 4° | $20 \pm 5^*$ | $0.21 \pm 0.12$ | $0.23 \pm 0.02$ |

\* means each conditions differed from the other with all $p < 0.05$ except EO versus Static ( $p = 0.75$ ).
$\beta$ means Amp 0.5° > Amp 1, 2 and 4° with $p < 0.05$ .
$\alpha$ means Amp 1° > Amp 2 and 4° with $p < 0.01$ .
$\mu$ means Amp 2° > Amp 4° with $p < 0.01$ .

## 3. Results and discussion

No difference was found between the EO and Static conditions in the mean FFT (Table 1). This suggests that wearing the VR headset did not alter baseline postural sway.

Each of the four VS amplitudes induced postural responses different from natural postural sway, as evidenced by increasing mean FFT and coherence spectra above the significance level at the tested frequencies (Figure 1 and Table 1). These results replicate previous findings from our group in another group of participants, and support that multi-sine VS induces postural responses within the 0.1-0.8 Hz frequency band in a VR-based paradigm.

**Figure 1.**
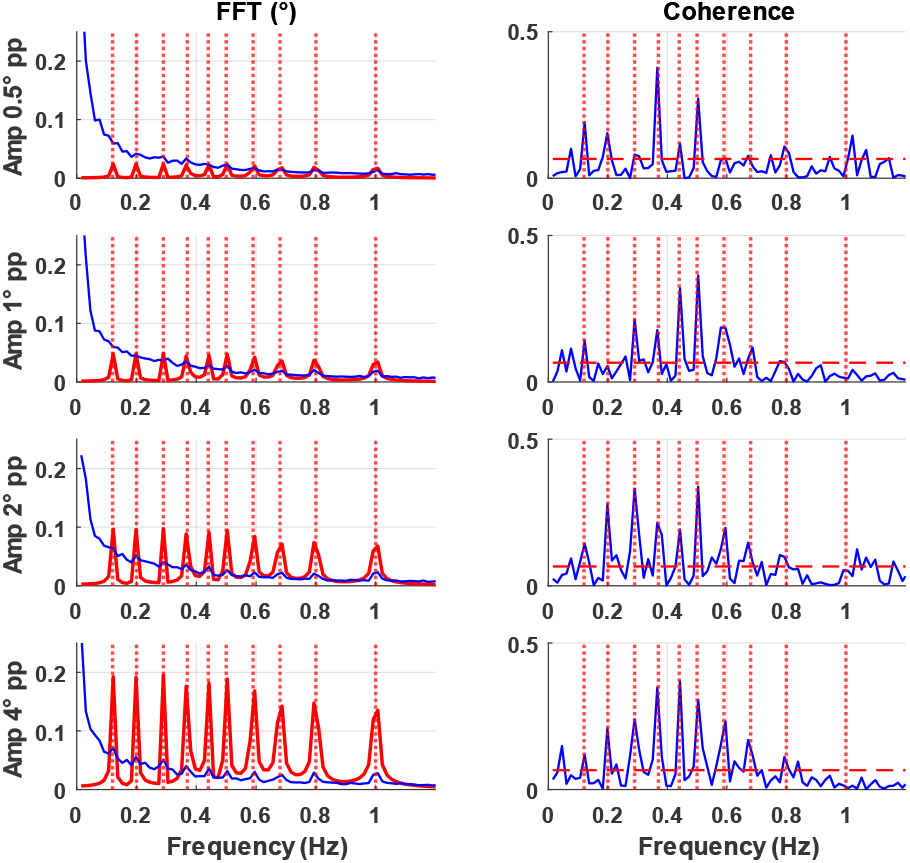
Mean group results of the frequency analysis. The left side indicates the FFT of the VS (red) and the AP body angle (blue). The right side shows the coherence between the VS and AP body angle. Horizontal red dashed lines indicate significance level of coherence (0.07), calculated for p < 0.05 based on the total length of the pooled signal. Credit: Baptiste Toussaint--Malard

When comparing the four dynamic conditions, FFT and coherence increased, while gain decreased with the VS amplitude (Table 1), replicating previous results (Assländer et al., 2023). These results support that the non-linearity of postural oscillations does not increase proportionally to stimulation amplitude Peterka (2002). Significant differences were found between the mean levels of the FFT and gain but not of the coherence across the four tested amplitudes (Table 1). Coherence visually peaked at 0.44 and/or 0.5 Hz (Figure 1), which is also consistent with previous reports (Peterka, 2002) suggesting greater postural responsiveness to intermediate-frequencies in the tested frequency range of visual inputs.

## 4. Conclusions

Multi-sine VS presented in a VR environment induces measurable postural responses within a specified frequency bandwidth (0.1 – 0.8 Hz). Increasing visual stimulus amplitude significatively increased postural oscillation, and tended to increase coherence but decreased gain. Among the four tested amplitudes, 2° was the lowest amplitude that still provides significant coherence at all stimulated frequencies (except for 1 Hz). These findings suggest that this amplitude may be suitable to study the contribution of visual inputs to postural control during movement execution in this specific frequency bandwidth.

## Acknowledgements

Authors thank participants and staff involved in this project.

## Conflict of Interest Statement

None

## Contributor Roles

BTM : Conceptualization, Data curation, Formal analysis, Investigation, Methodology, Software, Validation, Visualization, Writing-original draft, Writing-review & editing. FD : Conceptualization, Investigation, Methodology, Supervision, Validation, Visualization, Writing-original draft, Writing-review & editing. CLM : Investigation, Validation, Writing-review & editing. AD : Investigation, Validation, Writing-review & editing. PL : Investigation, Validation, Writing-review & editing. MB : Investigation, Validation, Writing-review & editing. RT : Conceptualization, Funding acquisition, Investigation, Methodology, Project administration, Resources, Supervision, Validation, Visualization, Writing-original draft, Writing-review & editing.

Based on CrediT: https://credit.niso.org/ :

## Funding

This project was funded by the Agence Nationale de la Recherche (ANR) project ANR-24-CE28-4602-01.

## Data, software, code availability

The raw data supporting the conclusions of this study will be made available on reasonable request.

## Notes

### Competing Interest Statement

The authors have declared no competing interest.

